# Simulating the outcome of amyloid treatments in Alzheimer’s disease from imaging and clinical data

**DOI:** 10.1101/2020.09.02.279521

**Authors:** Clément Abi Nader, Nicholas Ayache, Giovanni B. Frisoni, Philippe Robert, Marco Lorenzi, for the Alzheimer’s Disease Neuroimaging Initiative

**Affiliations:** Université Côte d’Azur, INRIA Sophia Antipolis, EPIONE Research Project, France; Memory Clinic and LANVIE-Laboratory of Neuroimaging of Aging, Hospitals and University of Geneva, Geneva, Switzerland; Université Côte d’Azur, CoBTeK lab, MNC3 program, France

**Keywords:** Alzheimer’s Disease, Clinical trials, Disease progression, Amyloid hypothesis, Biomarkers

## Abstract

In this study we investigate a novel quantitative instrument for the development of intervention strategies for disease modifying drugs in Alzheimer’s disease. Our framework is based on the modeling of the spatio-temporal dynamics governing the joint evolution of imaging and clinical biomarkers along the history of the disease, and allows the simulation of the effect of intervention time and drug dosage on the biomarkers’ progression. When applied to multi-modal imaging and clinical data from the Alzheimer’s Disease Neuroimaging Initiative our method enables to generate hypothetical scenarios of amyloid lowering interventions. The results quantify the crucial role of intervention time, and provide a theoretical justification for testing amyloid modifying drugs in the pre-clinical stage. Our experimental simulations are compatible with the outcomes observed in past clinical trials, and suggest that anti-amyloid treatments should be administered at least 7 years earlier than what is currently being done in order to obtain statistically powered improvement of clinical endpoints.

## Introduction

The number of people affected by Alzheimer’s disease has recently exceeded 46 millions and is expected to double every 20 years (Prince et al., 2015), thus posing significant healthcare challenges. Yet, while the disease mechanisms remain in large part unknown, there are still no effective pharmacological treatments leading to tangible improvements of patients’ clinical progression. One of the main challenges in understanding Alzheimer’s disease is that its progression goes through a silent asymptomatic phase that can stretch over decades before a clinical diagnosis can be established based on cognitive and behavioral symptoms. To help designing appropriate intervention strategies, hypothetical models of the disease history have been proposed, characterizing the progression by a cascade of morphological and molecular changes affecting the brain, ultimately leading to cognitive impairment (Jack et al., 2013; Jack & Holtzman, 2013). The dominant hypothesis is that disease dynamics along the asymptomatic period are driven by the deposition in the brain of the amyloid *β*peptide, triggering the so-called “amyloid cascade” (Bateman et al., 2012; Braak & Braak, 1991; Delacourte et al., 1999; Murphy & LeVine, 2010; Villemagne et al., 2013). Based on this rationale, clinical trials have been focusing on the development and testing of disease modifiers targeting amyloid *β* aggregates (Cummings, Lee, et al., 2019), for example by increasing its clearance or blocking its accumulation. Although the amyloid hypothesis has been recently invigorated by a post-hoc analysis of the aducanumab trial (Howard & Liu, 2020), clinical trials failed so far to show efficacy of this kind of treatments (Schwarz et al., 2019), as the clinical primary endpoints were not met (Egan et al., 2019; Honig et al., 2018; Wessels et al., 2019), or because of unacceptable adverse effects (Henley et al., 2019). In the past years, growing consensus emerged about the critical importance of intervention time, and about the need of starting anti-amyloid treatments during the pre-symptomatic stages of the disease (Aisen et al., 2018). Nevertheless, the design of optimal intervention strategies is currently not supported by quantitative analysis methods allowing to model and assess the effect of intervention time and dosing (Klein et al., 2019). The availability of models of the pathophysiology of Alzheimer’s disease would entail great potential to test and analyze clinical hypothesis characterizing Alzheimer’s disease mechanisms, progression, and intervention scenarios.

Within this context, quantitative models of disease progression, Disease progression Models referred to as DPMs, have been proposed (Fonteijn et al., 2012; Jedynak et al., 2012; Nader et al., 2020; Oxtoby et al., 2017; Schiratti et al., 2015), to quantify the dynamics of the changes affecting the brain during the whole disease span. These models rely on the statistical analysis of large datasets of different data modalities, such as clinical scores, or brain imaging measures derived from MRI, Amyloid-and Fluorodeoxyglucose-PET (Bilgel et al., 2015; Burnham et al., 2020; Donohue et al., 2014; Y Iturria-Medina et al., 2016; Koval et al., 2018). In general, DPMs estimate a long-term disease evolution from the joint analysis of multivariate time-series acquired on a short-term time-scale. Due to the temporal delay between the disease onset and the appearance of the first symptoms, DPMs rely on the identification of an appropriate temporal reference to describe the long-term disease evolution (Lorenzi et al., 2017; Marinescu et al., 2019). These tools are promising approaches for the analysis of clinical trials data, as they allow to represent the longitudinal evolution of multiple biomarkers through a global model of disease progression. Such a model can be subsequently used as a reference in order to stage subjects and quantify their relative progression speed (Insel et al., 2020; Li et al., 2019; Oxtoby et al., 2018; Young et al., 2014). However, these approaches remain purely descriptive as they don’t account for causal relationships among biomarkers. Therefore, they generally don’t allow to simulate progression scenarios based on hypothetical intervention strategies, thus providing a limited interpretation of the pathological dynamics. This latter capability is of utmost importance for planning and assessment of disease modifying treatments.

To fill this gap, recent works such as (Hao & Friedman, 2016; Petrella et al., 2019) proposed to model Alzheimer’s disease progression based on specific assumptions on the biochemical processes of pathological protein propagation. These approaches explicitly define biomarkers interactions through the specification of sets of Ordinary Differential Equations (ODEs), and are ideally suited to simulate the effect of drug interventions (Yasser Iturria-Medina et al., 2017). However, these methods are mostly based on the arbitrary choices of pre-defined evolution models, which are not inferred from data. This issue was recently addressed by (Garbarino & Lorenzi, 2019), where the authors proposed an hybrid modeling method combining traditional DPMs with dynamical models of Alzheimer’s disease progression. Still, since this approach requires to design suitable models of protein propagation across brain regions, extending this method to jointly account for spatio-temporal interactions between several processes, such as amyloid propagation, glucose metabolism, and brain atrophy, is considerably more complex. Finally, these methods are usually designed to account for imaging data only, which prevents to jointly simulate heterogeneous measures (Antelmi et al., 2019), such as image-based biomarkers and clinical outcomes, the latter remaining the reference markers for patients and clinicians.

In this work we present a novel computational model of Alzheimer’s disease progression allowing to simulate intervention strategies across the history of the disease. The model is here used to quantify the potential effect of amyloid modifiers on the progression of brain atrophy, glucose metabolism, and ultimately on the clinical outcomes for different scenarios of intervention. To this end, we model the joint spatio-temporal variation of different modalities along the history of Alzheimer’s disease by identifying a system of ODEs governing the pathological progression. This latent ODEs system is specified within an interpretable low-dimensional space relating multi-modal information, and combines clinically-inspired constraints with unknown interactions that we wish to estimate. The interpretability of the relationships in the latent space is ensured by mapping each data modality to a specific latent coordinate. The model is formulated within a Bayesian framework, where the latent representation and dynamics are efficiently estimated through stochastic variational inference. To generate hypothetical scenarios of amyloid lowering interventions, we apply our approach to multi-modal imaging and clinical data from the Alzheimer’s Disease Neuroimaging Initiative (ADNI). Our results provide a meaningful quantification of different intervention strategies, compatible with findings previously reported in clinical studies. For example, we estimate that in a study with 100 individuals per arm, statistically powered improvement of clinical endpoints can be obtained by completely arresting amyloid accumulation at least 11 years before Alzheimer’s dementia. The minimum intervention time decreases to 7 years for studies based on 1000 individuals per arm.

## Materials and methods

In the following sections, healthy individuals will be denoted as NL stable, subjects with mild cognitive impairment as MCI stable, subjects diagnosed with Alzheimer’s dementia as AD. We define conversion as the change of diagnosis towards a more pathological state. Therefore, NL converters are subjects who were diagnosed as cognitively normal at baseline and whose diagnosis changed either in MCI or AD during their follow-up visits. MCI converters are subjects who were diagnosed as MCI at baseline and subsequently progressed to AD. Diagnosis was established using the DX column from the ADNIMERGE file (https://adni.bitbucket.io/index.html), which reflects the standard ADNI clinical assessment based on Wechsler Memory Scale, Mini-Mental State Examination, and Clinical Dementia Rating. Amyloid concentration and glucose metabolism are respectively measured by (18)F-florbetapir Amyloid (AV45)-PET and (18)F-fluorodeoxyglucose (FDG)-PET imaging. Cognitive and functional abilities are assessed by the following neuro-psychological tests: Alzheimer’s Disease Assessment Scale (ADAS11), Mini-Mental State Examination (MMSE), Functional Assessment Questionnaire (FAQ), Rey Auditory Verbal Learning Test (RAVLT) immediate, RAVLT learning, RAVLT forgetting, and Clinical Dementia Rating Scale Sum of Boxes (CDRSB).

### Study cohort and biomarkers’ changes across clinical groups

Our study is based on a cohort of 442 amyloid positive individuals composed of 71 NL stable subjects, 33 NL converters subjects, 131 subjects diagnosed with MCI, 105 MCI converters subjects, and 102 AD patients. Among the 131 MCI subjects, 78 were early MCI and 53 were late MCI. Concerning the group of MCI converters, 80 subjects were late MCI at baseline and 25 were early MCI. The term ``amyloid positive’’ refers to subjects whose amyloid level in the CSF was below the nominal cutoff of 192 pg/ml (Gamberger et al., 2017) either at baseline, or during any follow-up visit, and conversion to AD was determined using the last available follow-up information. This preliminary selection of patients aims at constituting a cohort of subjects for whom it is more likely to observe “Alzheimer’s pathological changes” (Jack et al., 2018). The length of follow-up varies between 0 and 16 years. Further information about the data are available on https://adni.bitbucket.io/reference/, while details on data acquisition and processing are provided in Section *Data acquisition and preprocessing*. We show in Table 1A socio-demographic information for the training cohort across the different clinical groups. Table 1B shows baseline values and annual rates of change across clinical groups for amyloid burden (average normalized AV45 uptake in frontal cortex, anterior cingulate, precuneus and parietal cortex), glucose metabolism (average normalized FDG uptake in frontal cortex, anterior cingulate, precuneus and parietal cortex), for hippocampal and medial temporal lobe volumes, and for the cognitive ability as measured by ADAS11. Compatibly with previously reported results (Cash et al., 2015; Schuff et al., 2009), we observe that while regional atrophy, glucose metabolism and cognition show increasing rate of change when moving from healthy to pathological conditions, the change of AV45 is maximum in NL stable, NL converters and MCI stable subjects. We also notice the increased magnitude of ADAS11 in AD as compared to the other clinical groups. Finally, we note that glucose metabolism and regional atrophy show comparable magnitudes of change.

**Table 1.**
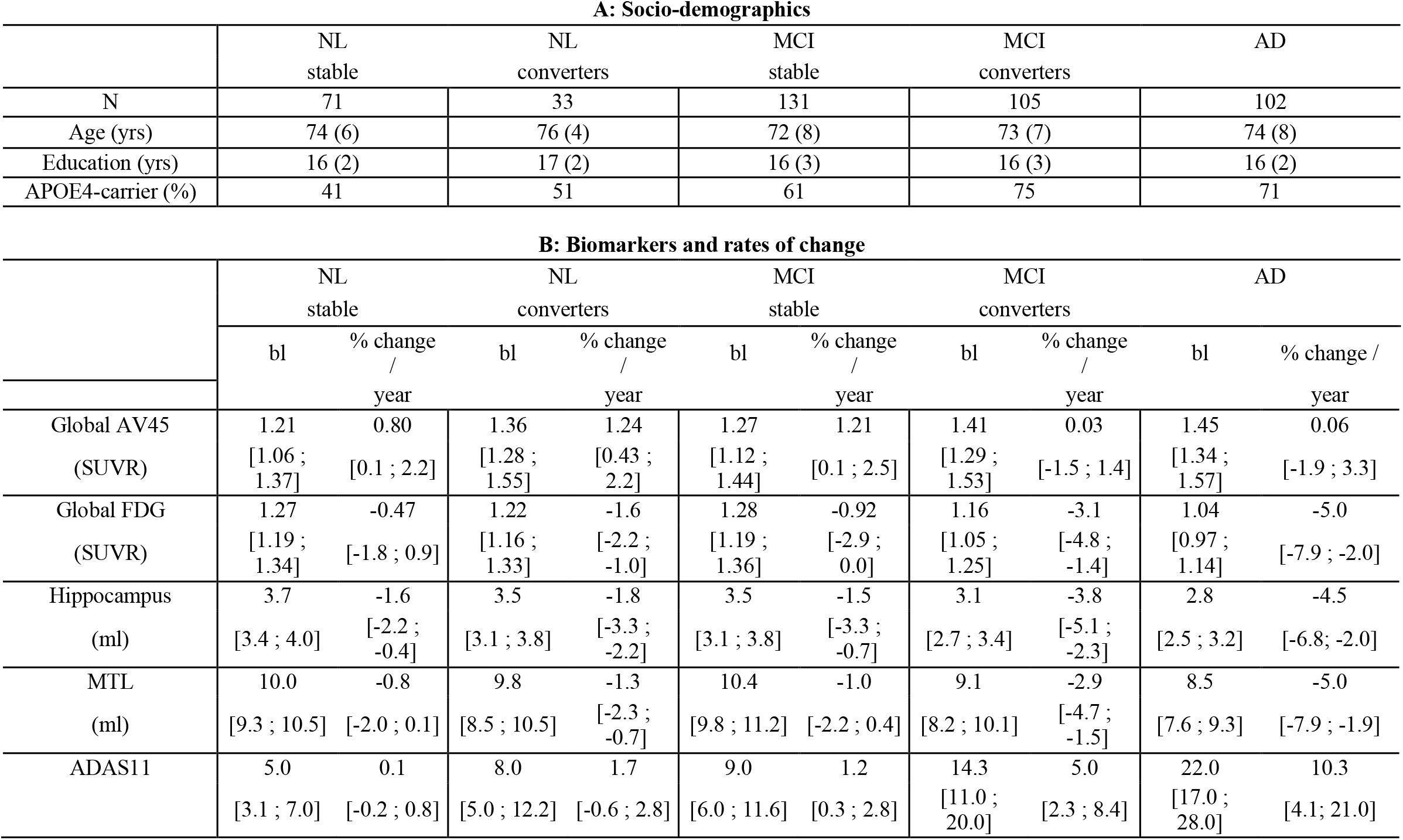
A: Baseline socio-demographic information for training cohort (442 subjects for 2781 data points, follow-up from 0 to 16 years depending on subjects). Average values, standard deviation in parenthesis. B: Baseline values (bl) and annual rates of change (\% change / year) of amyloid burden (average normalized AV45 uptake in frontal cortex, anterior cingulate, precuneus and parietal cortex), glucose metabolism (average normalized FDG uptake in frontal cortex, anterior cingulate, precuneus and parietal cortex), hippocampus volume, medial temporal lobe volume, and ADAS11 score for the different clinical groups. Median values, interquartile range below. The volumes of the hippocampus and the medial temporal lobe are averaged across left and right hemispheres. NL: healthy individuals, MCI: individuals with mild cognitive impairment, AD: patients with Alzheimer’s dementia. APOE4: apolipoprotein E ∊4. FDG: (18)F-fluorodeoxyglucose Positron Emission Tomography (PET) imaging. AV45: (18)F-florbetapir Amyloid PET imaging. SUVR: Standardized Uptake Value Ratio. MTL: Medial Temporal Lobe. ADAS11: Alzheimer’s Disease Assessment Scale-cognitive subscale, 11 items.

The observations presented in Table 1 provide us with a coarse representation of the biomarkers’ trajectories characterizing Alzheimer’s disease. The complexity of the dynamical changes we may infer is limited, as the clinical stages roughly approximate a temporal scale describing the disease history, while very little insights can be obtained about the biomarkers’ interactions. Within this context, our model allows the quantification of the fine-grained dynamical relationships across biomarkers at stake during the history of the disease. Investigation of intervention scenarios can be subsequently carried out by opportunely modulating the estimated dynamics parameters according to specific intervention hypothesis (e.g. amyloid lowering at a certain time).

### Model overview

We provide in Figure 1 an overview of the presented method. Baseline multi-modal imaging and clinical information for a given subject are transformed into a latent variable composed of four z-scores quantifying respectively the overall severity of atrophy, glucose metabolism, amyloid burden, and cognitive and functional assessment. The model estimates the dynamical relationships across these z-scores to optimally describe the temporal transitions between follow-up observations. These transition rules are here mathematically defined by the parameters of a system of ODEs, which is estimated from the data. This dynamical system allows to compute the evolution of the z-scores over time from any baseline observation, and to predict the associated multi-modal imaging and clinical measures. It is important to note that this modelling choice requires to have at least one visit per patient for which all the measures are available, in order to compute the z-scores temporal evolution.

**Figure 1.**
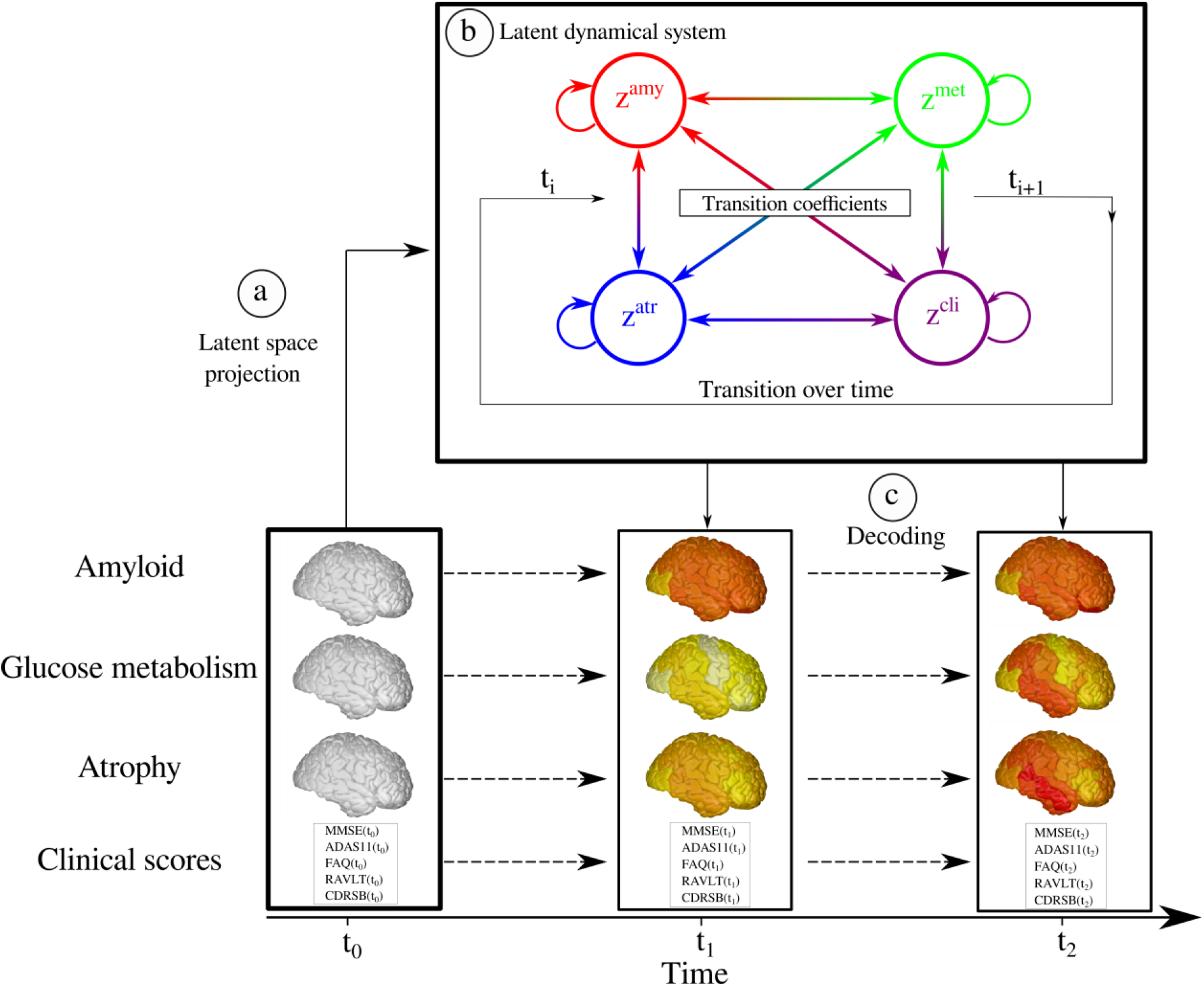
Overview of the method. a) High-dimensional multi-modal measures are projected into a 4-dimensional latent space. Each data modality is transformed in a corresponding z-score z^*amy*^, z^*met*^, z^*atr*^, z^*cli*^. b) The dynamical system describing the relationships between the z-scores allows to compute their transition across the evolution of the disease. c) Given the latent space and the estimated dynamics, the follow-up measurements can be reconstructed to match the observed data.

The model thus enables to simulate the pathological progression of biomarkers across the entire history of the disease. Once the model is estimated, we can modify the ODEs parameters to simulate different evolution scenarios according to specific hypothesis. For example, by reducing the parameters associated with the progression rate of amyloid, we can investigate the relative change in the evolution of the other biomarkers. This setup thus provides us with a data-driven system enabling the exploration of hypothetical intervention strategies, and their effect on the pathological cascade.

### Data modelling

We consider observations 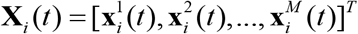, which correspond to multivariate measures derived from *M* different modalities (e.g clinical scores, MRI, AV45, or FDG measures) at time *t* for subject *i*. Each vector 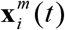 has dimension *D_m_*. We postulate the following generative model, in which the modalities are assumed to be independently generated by a common latent representation of the data **z**_*i*_ (*t*):

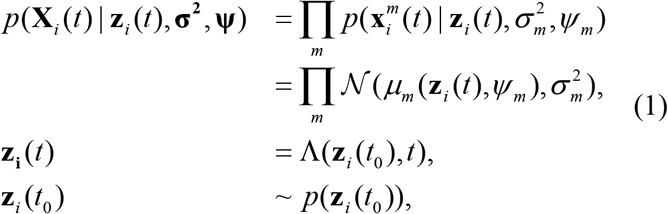

where 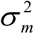 is measurement noise, while *ψ*_*m*_ are the parameters of the function *μ*_*m*_ which maps the latent state to the data space for the modality *m*. For simplicity of notation we denote **z**_*i*_ (*t*) by **z**(*t*). We assume that each coordinate of **z** is associated to a specific modality *m*, leading to an *M*-dimensional latent space. The ∧ operator which gives the value of the latent representation at a given time *t*, is defined by the solution of the following system of ODEs:

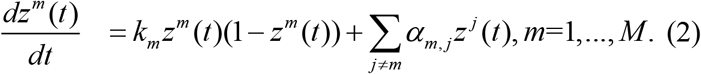

For each coordinate, the first term of the equation enforces a sigmoidal evolution with a progression rate *k_m_*, while the second term accounts for the relationship between modalities *m* and *j* through the parameters *α*_*m,j*_. This system can be rewritten as:

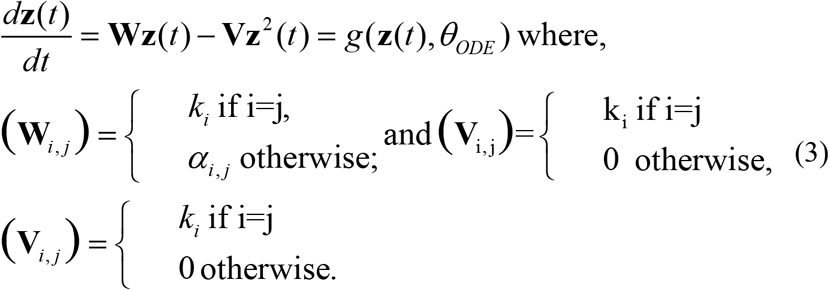

*θ_ODE_* denotes the parameters of the system of ODEs, which correspond to the entries of the matrices **W** and **V**. According to Equation (3), for each initial condition **z**(0), the latent state at time *t* can be computed through integration, 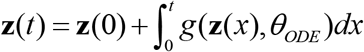.

We resort to variational inference and stochastic gradient descent in order to optimize the parameters of the model. The procedure is detailed in Sections *Variational inference* and *Model optimization* of the Supplementary Material.

### Simulating the long-term progression of Alzheimer’s disease

To simulate the long-term progression of Alzheimer’s disease we first project the AD subjects in the latent space via the encoding functions. We can subsequently follow the trajectories of these subjects backward and forward in time, in order to estimate the associated trajectory from the healthy to their respective pathological condition. In practice, a Gaussian Mixture Model is used to fit the empirical distribution of the AD subjects’ latent projection. The number of components and covariance type of the Gaussian Mixture Model is selected by relying on the Akaike information criterion (Akaike, 1998). The fitted Gaussian Mixture Model allows us to sample pathological latent representations **z**_*i*_ (*t*_0_) that can be integrated forward and backward in time thanks to the estimated set of latent ODEs, to finally obtain a collection of latent trajectories **Z**(*t*) = [**z**_1_(*t*),…,**z**_*N*_(*t*)] summarizing the distribution of the long-term Alzheimer’s disease evolution.

### Simulating intervention

In this section we assume that we computed the average latent progression of the disease **z**(*t*). Thanks to the modality-wise encoding (*cf*. Supplementary section *Variational inference*) each coordinate of the latent representation can be interpreted as representing a single data modality. Therefore, we propose to simulate the effect of a hypothetical intervention on the disease progression, by modulating the vector 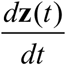 after each integration step such that:

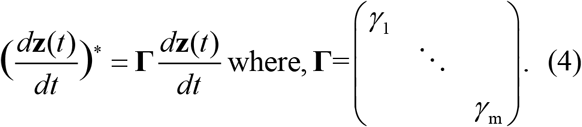

The values *γ*_*m*_ are fixed between 0 and 1, allowing to control the influence of the corresponding modalities on the system evolution, and to create hypothetical scenarios of evolution. For example, for a 100% (resp. 50%) amyloid lowering intervention we set *γ*_*amy*_ = 0(resp. *γ*_*amy*_ = 0.5).

### Evaluating disease severity

Given an evolution **z**(*t*) describing the disease progression in the latent space, we propose to consider this trajectory as a reference and to use it in order to quantify the individual disease severity of a subject **X**. This is done by estimating a time-shift *τ* defined as:

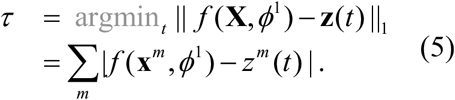

This time-shift allows to quantify the pathological stage of a subject with respect to the disease progression along the reference trajectory **z**(*t*). Moreover, the time-shift can still be estimated even in the case of missing data modalities, by only encoding the available measures of the observed subject.

### Statistical analysis

The model was implemented using the Pytorch library (Paszke et al., 2019). The estimated disease severity was compared group-wise via two-sided Wilcoxon-Mann-Whitney test (P < 0.01). Differences between the clinical outcomes distribution after simulation of intervention were compared via two-sided Student’s T-test (P < 0.01). Shadowed areas in the different figures show ± standard deviation of the mean.

### Data availability

The data used in this study are available from the ADNI database (adni.loni.usc.edu).

## Results

In the following, MRI, FDG-PET, and AV45-PET images are processed in order to respectively extract regional gray matter density, glucose metabolism and amyloid load from a brain parcellation. The z-scores of gray matter atrophy (z^atr^), glucose metabolism (z^met^), and amyloid burden (z^amy^), are computed using the measures obtained by this pre-processing step. The clinical z-score z^cli^ is derived from neuro-psychological scores: ADAS11, MMSE, FAQ, RAVLT immediate, RAVLT learning, RAVLT forgetting and CDRSB. This panel of scores was chosen to provide a comprehensive representation of cognitive, memory and functional abilities.

### Data acquisition and preprocessing

Data used in the preparation of this article were obtained from the ADNI database. The ADNI was launched in 2003 as a public-private partnership, led by Principal Investigator Michael W. Weiner, MD. For up-to-date information, see www.adni-info.org.

We considered four types of biomarkers, related to clinical scores, gray matter atrophy, amyloid load and glucose metabolism, and respectively denoted by *cli*, *atr*, *amy* and *met*. MRI images were processed following the longitudinal pipeline of Freesurfer (Reuter et al., 2012), to obtain gray matter volumes in a standard anatomical space. AV45-PET and FDG-PET images were aligned to the closest MRI in time and normalized to the cerebellum uptake. Regional gray matter density, amyloid load and glucose metabolism were extracted from the Desikan-Killiany parcellation (Desikan et al., 2006). We discarded white-matter, ventricular, and cerebellar regions, thus obtaining 82 regions that were averaged across hemispheres. Therefore, for a given subject, **x**^*atr*^, **x**^*amy*^ and **x**^*met*^ are respectively 41-dimensional vectors. The variable **x**^*cli*^ is composed of the neuro-psychological scores ADAS11, MMSE, RAVLT immediate, RAVLT learning, RAVLT forgetting, FAQ, and CDRSB. The total number of measures is of 2781 longitudinal data points. We recall that the model estimation requires a visit for which all the measures are available in order to obtain the z-scores evolution of a given subject, but can handle missing data in the follow-up by finding the parameters that best match the available measures.

### Progression model and latent relationships

We show in Figure 2 panel I) the dynamical relationships across the different z-scores estimated by the model, where direction and intensity of the arrows quantify the estimated increase of one variable with respect to the other. Being the scores adimensional, they have been conveniently rescaled to the range [0,1] indicating increasing pathological levels. These relationships extend the summary statistics reported in Table 1 to a much finer temporal scale and wider range of possible biomarkers’ values. We observe in Figure 2A, 2B and 2C that large values of the amyloid score z^*amy*^ trigger the increase of the remaining ones: z^*met*^, z^*atr*^, and z^*cli*^. Figure 2D shows that large increase of the atrophy score z^*atr*^ is associated to pathological glucose metabolism indicated by large values of z^*met*^. Moreover, we note that high z^*met*^ values also contribute to an increase of z^*cli*^ (Figure 2E). Finally, Figure 2F shows that high atrophy values lead to an increase mostly along the clinical dimension z^*cli*^. This chain of relationships is in agreement with the cascade hypothesis of AD (Jack et al., 2013; Jack & Holtzman, 2013).

**Figure 2.**
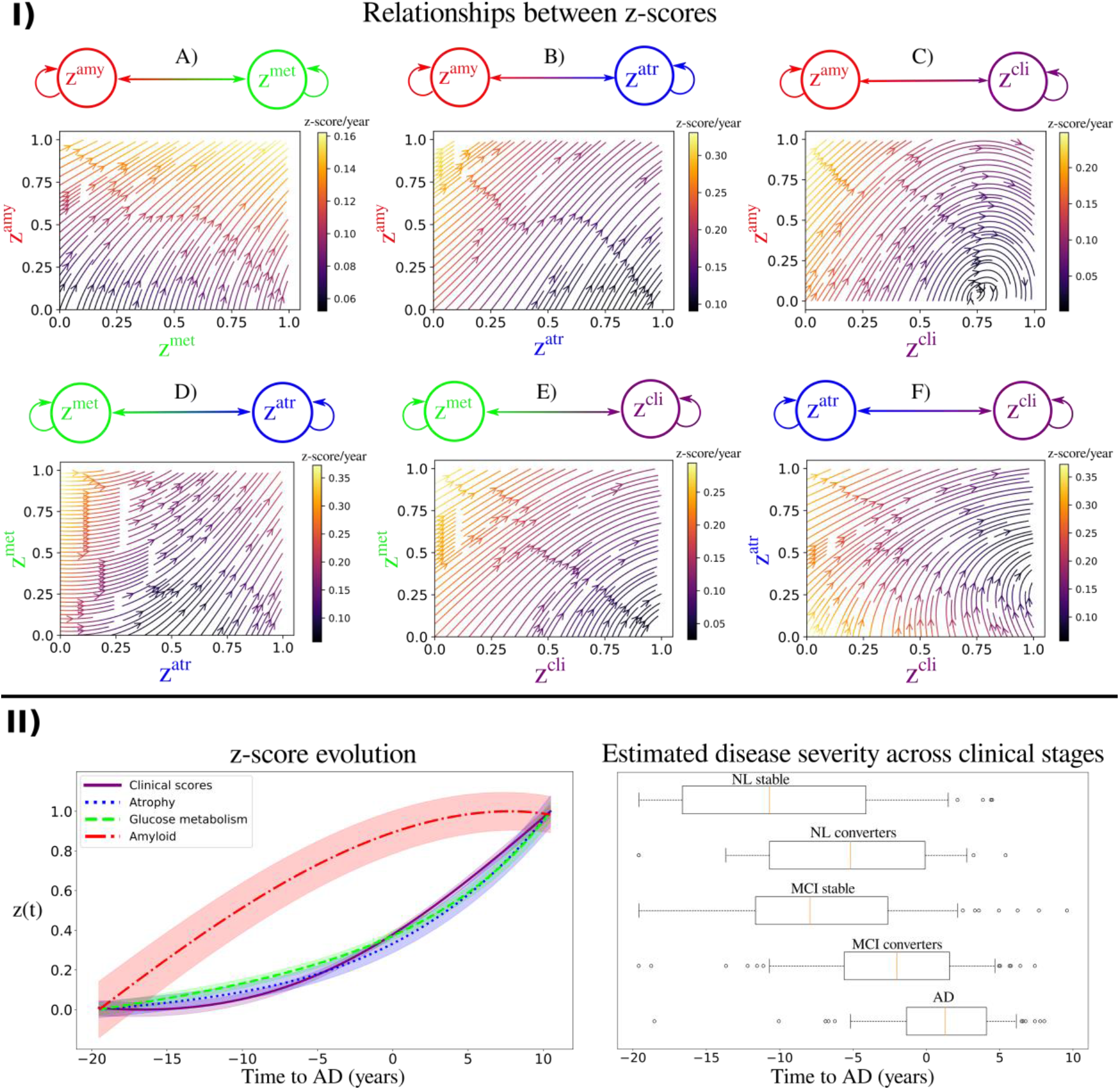
Dynamical relationships, z-scores evolution and disease staging. Panel I: Estimated dynamical relationships across the different z-scores (A to F). Given the values of two z-scores, the arrow at the corresponding coordinates indicates how one score evolves with respect to the other. The intensity of the arrow gives the strength of the relationship between the two scores. Panel II, left: Estimated long-term latent dynamics (time is relative to conversion to Alzheimer’s dementia). Shadowed areas represent the standard deviation of the average trajectory. Panel II, right: Distribution of the estimated disease severity across clinical stages, relatively to the long-term dynamics on the left. NL: normal individuals, MCI: mild cognitive impairment, AD: Alzheimer’s dementia.

Relying on the dynamical relationships shown in Figure 2 panel I), starting from any initial set of biomarkers values we can estimate the relative trajectories over time. Figure 2 panel II) (left), shows the evolution obtained by extrapolating backward and forward in time the trajectory associated to the z-scores of the AD group. The x-axis represents the years from conversion to AD, where the instant *t*=0 corresponds to the average time of diagnosis estimated for the group of MCI progressing to dementia. As observed in Figure 2 panel I) and Table 1, the amyloid score z^*amy*^ increases and saturates first, followed by z^*met*^ and z^*atr*^ scores whose progression slows down when reaching clinical conversion, while the clinical score exhibits strong acceleration in the latest progression stages. Figure 2 panel II) (right) shows the group-wise distribution of the disease severity estimated for each subject relatively to the modelled long-term latent trajectories. The group-wise difference of disease severity across groups is statistically significant and increases when going from healthy to pathological stages (Wilcoxon-Mann-Whitney test p < 0.01 for each comparisons). The reliability of the estimation of disease severity was further assessed through testing on an independent cohort, and by comparison with a previously proposed disease progression modeling method from the state-of-the-art (Lorenzi et al., 2017). The results are provided in section *Time-shift comparison* and *validation* of the Supplementary Material and show positive generalization results as well as a favorable comparison with the benchmark method.

From the z-score trajectories of Figure 2 panel II) (left) we predict the progression of imaging and clinical measures shown in Figure 3. We observe that amyloid load globally increases and saturates early, compatibly with the positive amyloid condition of the study cohort. Abnormal glucose metabolism and gray matter atrophy are delayed with respect to amyloid, and tend to map prevalently temporal and parietal regions. Finally, the clinical measures exhibit a non-linear pattern of change, accelerating during the latest progression stages. These dynamics are compatible with the summary measures on the raw data reported in Table 1.

**Figure 3.**
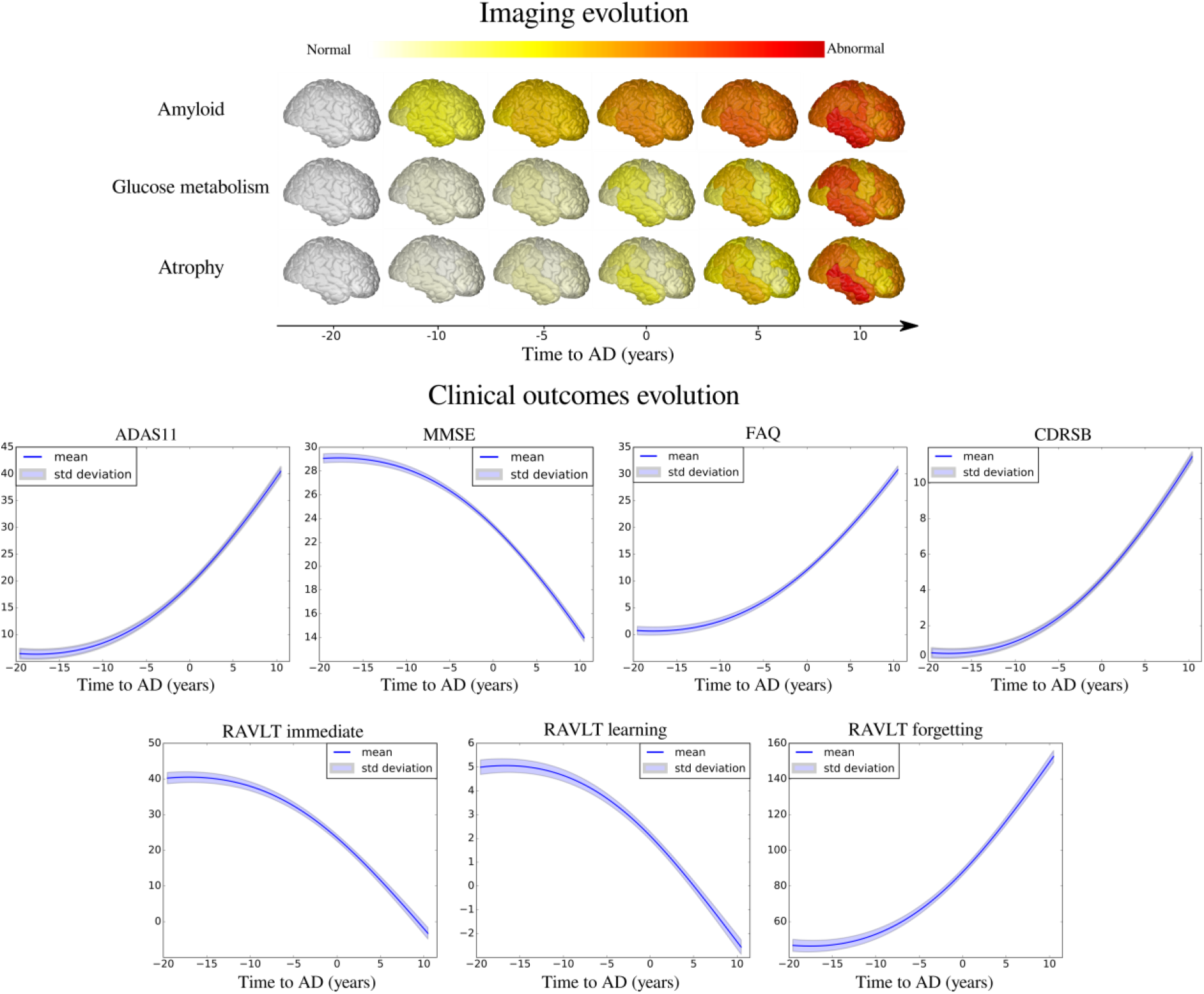
Model-based progression of Alzheimer’s disease. Estimated long-term evolution of cortical measurements for the different types of imaging markers, and clinical scores. Shadowed areas represent the standard deviation of the average trajectory. Brain images were generated using the software provided in (Marinescu et al., 2019).

### Simulating clinical intervention

This experimental section is based on two intervention scenarios: a first one in which amyloid is lowered by 100%, and a second one in which it is reduced by 50% with respect to the estimated natural progression. In Figure 4 we show the latent z-scores evolution resulting from either 100% or 50% amyloid lowering performed at the time *t*=-20 years. According to these scenarios, intervention results in a sensitive reduction of the pathological progression for atrophy, glucose metabolism and clinical scores, albeit with a stronger effect in case of total blockage.

**Figure 4.**
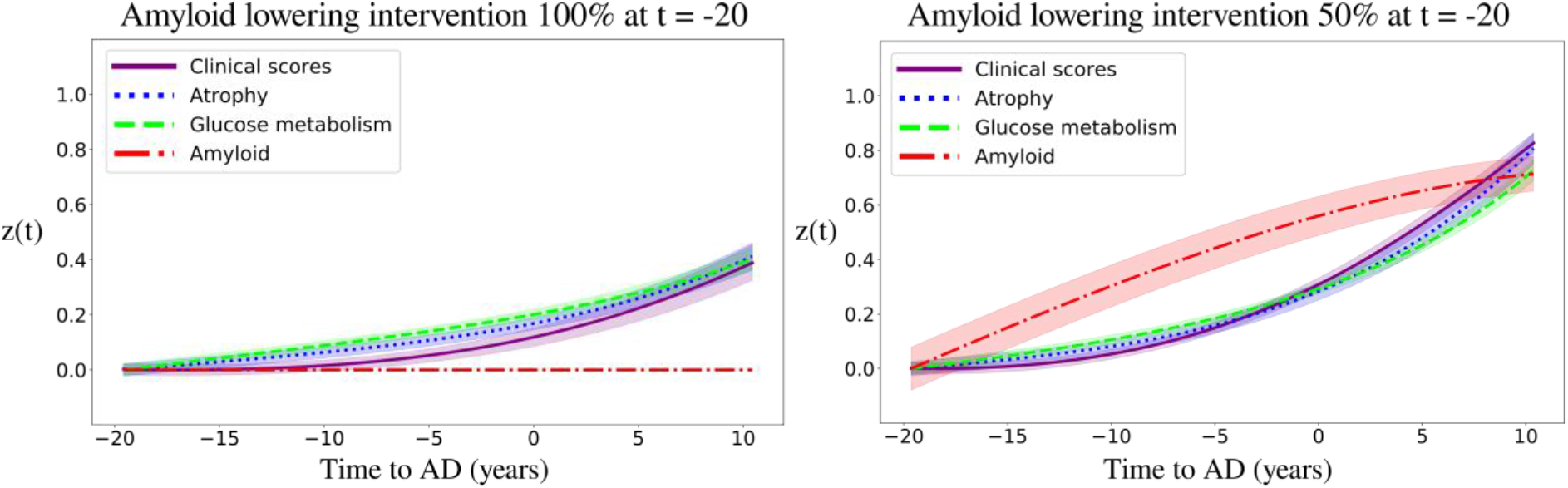
Simulation of amyloid lowering intervention on the z-scores evolution. Hypothetical scenarios of irreversible amyloid lowering interventions at *t*=-20 years from Alzheimer’s dementia diagnosis, with a rate of 100 % (left) or 50% (right). Shadowed areas represent the standard deviation of the average trajectory.

We further estimated the resulting clinical endpoints associated with the two amyloid lowering scenarios, at increasing time points and for different sample sizes. Clinical endpoints consisted in the simulated ADAS11, MMSE, FAQ, RAVLT immediate, RAVLT learning, RAVLT forgetting and CDRSB scores at the reference conversion time (*t*=0). The case placebo indicates the scenario where clinical values were computed at conversion time from the estimated natural progression shown in Figure 2 panel II) (left). Figure 5 shows the change in statistical power depending on intervention time and sample sizes. For large sample sizes (1000 subjects per arm) a power greater than 0.8 can be obtained around 7 years before conversion, depending on the outcome score, where in general we observe that RAVLT forgetting exhibits a higher power than the other scores. When sample size is lower than 100 subjects per arm, a power greater than 0.8 is reached if intervention is performed at the latest 11 years before conversion, with a mild variability depending on the considered clinical score. We notice that in the case of 50% amyloid lowering, in order to reach the same power intervention needs to be consistently performed earlier compared to the scenario of 100% amyloid lowering for the same sample size and clinical score. For instance, if we consider ADAS11 with a sample size of 100 subjects per arm, a power of 0.8 is obtained for a 100% amyloid lowering intervention performed 11.5 years before conversion, while in case of a 50% amyloid lowering the equivalent effect would be obtained by intervening 15 years before conversion.

**Figure 5.**
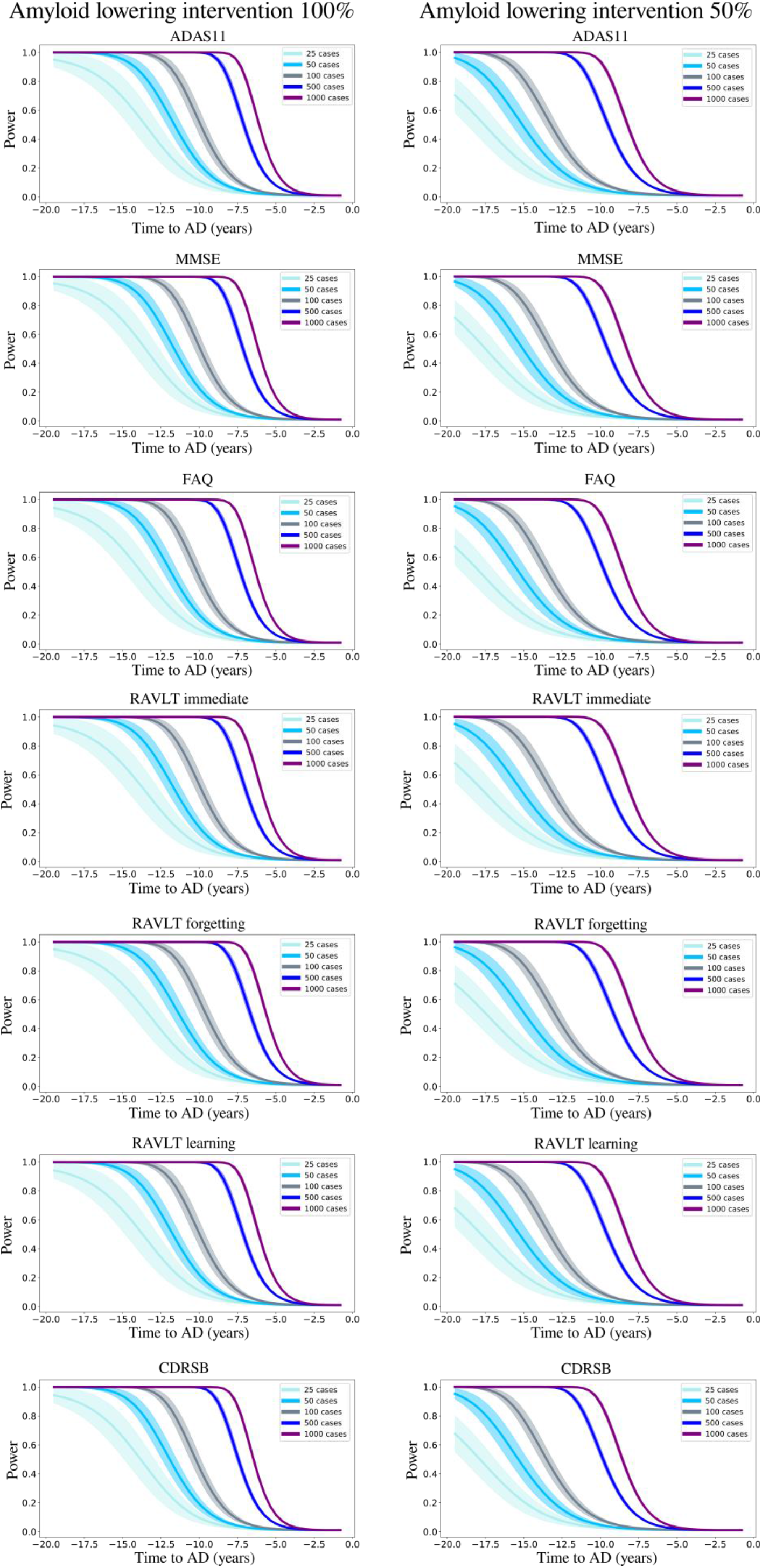
Evolution of the statistical power in different intervention scenarios. Statistical power of the Student t-test comparing the estimated clinical outcomes at conversion time between placebo and treated scenarios, according to the year of simulated intervention (100% and 50% amyloid lowering) and sample size.

We provide in Table 2 the estimated improvement for each clinical score at conversion with a sample size of 100 subjects per arm for both 100% and 50% amyloid lowering depending on the intervention time. We observe that for the same intervention time, 100% amyloid lowering always results in a larger improvement of clinical endpoints compared to 50% amyloid lowering. We also note that in the case of 100% lowering, clinical endpoints obtained for intervention at *t*=-15 years correspond to typical cutoff values for inclusion into Alzheimer’s disease trials (ADAS11 = 13.7 ± 5.8, MMSE = 25.7± 2.5, see Supplementary Table 2) (Gamberger et al., 2017; Kochhann et al., 2010).

**Table 2:** Estimated mean (standard deviation) improvement of clinical outcomes at predicted conversion time for the normal progression case by year of simulated intervention (100% and 50% amyloid lowering interventions). Results in bold indicate a statistically significant difference between placebo and treated scenarios (p<0.01, two-sided t-test, 100 cases per arm). AD: Alzheimer’s dementia, ADAS11: Alzheimer’s Disease Assessment Scale, MMSE: Mini-Mental State Examination, FAQ: Functional Assessment Questionnaire, RAVLT: Rey Auditory Verbal Learning Test, CDRSB: Clinical Dementia rating Scale Sum of Boxes.

|  | -20 | -15 | -12.5 | -10 | -5 | -3 | -2 | -1 |
| --- | --- | --- | --- | --- | --- | --- | --- | --- |
| ADAS11 | **11.1, (6.4)** | **5.2, (2.9)** | **3.0, (1.7)** | **1.6, (1.0)** | 0.3, (0.2) | 0.1, (0.1) | 0.0, (0.0) | 0.0, (0.0) |
| MMSE | **4.9, (2.8)** | **2.3, (1.3)** | **1.3, (0.8)** | **0.7, (0.4)** | 0.1, (0.1) | 0.0, (0.0) | 0.0, (0.0) | 0.0, (0.0) |
| FAQ | **9.6, (5.6)** | **4.5, (2.5)** | **2.6, (1.5)** | **1.4, (0.8)** | 0.2, (0.2) | 0.1, (0.1) | 0.0, (0.0) | 0.0, (0.0) |
| RAVLT immediate | **15.3, (8.9)** | **7.2, (4.1)** | **4.2, (2.4)** | **2.3, (1.4)** | 0.5, (0.3) | 0.2, (0.1) | 0.1, (0.1) | 0.0, (0.0) |
| RAVLT learning | **2.7, (1.6)** | **1.3, (0.7)** | **0.7, (0.4)** | **0.4, (0.2)** | 0.1, (0.1) | 0.0, (0.0) | 0.0, (0.0) | 0.0, (0.0) |
| RAVLT forgetting | **37.2, (21.5)** | **17.7, (9.9)** | **10.5, (6.0)** | **5.8, (3.5)** | 1.3, (0.9) | 0.5, (0.4) | 0.2, (0.2) | 0.1, (0.1) |
| CDRSB | **3.5, (2.0)** | **1.6, (0.9)** | **0.9, (0.5)** | **0.5, (0.3)** | 0.1, (0.1) | 0.0, (0.0) | 0.0, (0.0) | 0.0, (0.0) |

**Amyloid lowering intervention 50%** Point improvement per intervention time
|  | -20 | -15 | -12.5 | -10 | -5 | -3 | -2 | -1 |
| --- | --- | --- | --- | --- | --- | --- | --- | --- |
| ADAS11 | **5.0, (2.5)** | **2.4, (1.2)** | **1.4, (0.7)** | 0.8, (0.4) | 0.2, (0.1) | 0.1, (0.0) | 0.0, (0.0) | 0.0, (0.0) |
| MMSE | **2.2, (1.1)** | **1.0, (0.5)** | **0.6, (0.3)** | 0.4, (0.2) | 0.1, (0.0) | 0.0, (0.0) | 0.0, (0.0) | 0.0, (0.0) |
| FAQ | **4.3, (2.1)** | **2.0, (1.0)** | **1.2, (0.6)** | 0.7, (0.4) | 0.1, (0.1) | 0.0, (0.0) | 0.0, (0.0) | 0.0, (0.0) |
| RAVLT immediate | **6.9, (3.4)** | **3.3, (1.6)** | **1.9, (1.0)** | 1.2, (0.6) | 0.2, (0.1) | 0.1, (0.1) | 0.0, (0.0) | 0.0, (0.0) |
| RAVLT learning | **1.2, (0.6)** | **0.6, (0.3)** | **0.3, (0.2)** | 0.2, (0.1) | 0.0, (0.0) | 0.0, (0.0) | 0.0, (0.0) | 0.0, (0.0) |
| RAVLT forgetting | **16.7, (8.2)** | **8.1, (4.0)** | **4.8, (2.5)** | 2.9, (1.6) | 0.6, (0.4) | 0.2, (0.2) | 0.1, (0.1) | 0.0, (0.0) |
| CDRSB | **1.6, (0.8)** | **0.7, (0.4)** | **0.4, (0.2)** | 0.2, (0.1) | 0.0, (0.0) | 0.0, (0.0) | 0.0, (0.0) | 0.0, (0.0) |

## Discussion

We presented a framework to jointly model the progression of multi-modal imaging and clinical data, based on the estimation of latent biomarkers’ relationships governing Alzheimer’s disease progression. The model is designed to simulate intervention scenarios in clinical trials, and in this study we focused on assessing the effect of anti-amyloid drugs on biomarkers’ evolution, by quantifying the effect of intervention time and drug efficacy on clinical outcomes. Our results underline the critical importance of intervention time, which should be performed sensibly early during the pathological history to effectively appreciate the effectiveness of disease modifiers.

The results obtained with our model are compatible with findings reported in recent clinical studies (Egan et al., 2019; Honig et al., 2018; Wessels et al., 2019). For example, if we consider 500 patients per arm and perform a 100% amyloid lowering intervention for 2 years to reproduce the conditions of the recent trial of Verubecestat (Egan et al., 2019), the average improvement of MMSE predicted by our model is of 0.02, falling in the 95% confidence interval measured during that study ([−0.5; 0.8]). While recent anti-amyloid trials such as (Egan et al., 2019; Honig et al., 2018; Wessels et al., 2019) included between 500 and 1000 mild AD subjects per arm and were conducted over a period of two years at most, our analysis suggests that clinical trials performed with less than 1000 subjects with mild AD may be consistently under-powered. Indeed, we see in Figure 5 that with a sample size of 1000 subjects per arm and a total blockage of amyloid production, a power of 0.8 can be obtained only if intervention is performed at least 7 years before conversion.

These results allow to quantify the crucial role of intervention time, and provide a theoretical justification for testing amyloid modifying drugs in the pre-clinical stage (Aisen et al., 2018; Sperling et al., 2011). This is for example illustrated in Table 2, in which we notice that clinical endpoints are close to placebo even when the simulated intervention takes place 10 years before conversion, while stronger cognitive and functional changes happen when amyloid is lowered by 100% or 50% earlier. These findings may be explained by considering that amyloid accumulates over more than a decade, and that when amyloid clearance occurs the pathological cascade is already entrenched (Rowe et al., 2010). Our results are thus supporting the need to identify subjects at the pre-clinical stage, that is to say still cognitively normal, which is a challenging task. Currently, one of the main criteria to enroll subjects into clinical trials is the presence of amyloid in the brain, and blood-based markers are considered as potential candidates for identifying patients at risk for Alzheimer’s disease (Zetterberg & Burnham, 2019). Moreover, recent works such as (Blennow et al., 2010; Westwood et al., 2016) have proposed more complex entry criteria to constitute cohorts based on multi-modal measurements. Within this context, our model could also be used as an enrichment tool by quantifying the disease severity based on multi-modal data as shown in Figure 2 panel II) (right). Similarly, the method could be applied to predict the evolution of single patient given its current available measurements.

An additional critical aspect of anti-amyloid trials is the effect of dose exposure on the production of amyloid (Klein et al., 2019). Currently, *β*-site amyloid precursor protein cleaving enzyme (BACE) inhibitors allow to suppress amyloid production from 50% to 90%. In this study we showed that lowering amyloid by 50% consistently decreases the treatment effect compared to a 100% lowering at the same time. For instance, if we consider a sample size of 1000 subjects per arm in the case of a 50% amyloid lowering intervention, 80% power can be reached only 10 years before conversion instead of 7 years for a 100% amyloid lowering intervention. This ability of our model to control the rate of amyloid progression is fundamental in order to provide realistic simulations of anti-amyloid trials.

In Figure 2 panel I) we showed that amyloid triggers the pathological cascade affecting the other markers, thus confirming its dominating role on disease progression. Assuming that the data used to estimate the model is sufficient to completely depict the history of the pathology, our model can be interpreted from a causal perspective. However, we cannot exclude the existence of other mechanisms driving amyloid accumulation, which our model cannot infer from the existing data. Therefore, our findings should be considered with care, while the integration of additional biomarkers of interest will be necessary to account for multiple drivers of the disease. It is worth noting that recent works ventured the idea to combine drugs targeting multiple mechanisms at the same time (Gauthier et al., 2019). For instance, pathologists have shown tau deposition in brainstem nuclei in adolescents and children (Kaufman et al., 2018), and clinicians are currently investigating the pathological effect of early tau spreading on Alzheimer’s disease progression (Pontecorvo et al., 2019), raising crucial questions about its relationship with amyloid accumulation, and the impact on cognitive impairment (Cummings, Blennow, et al., 2019). In this study, 190 subjects underwent at least one Tau-PET scan. However, when considering the subjects for whom there exists one visit in which all the data modalities were available, the number of patients in the study cohort decreased to 33. This low sample size prevented us from estimating reliable trajectories for this biomarker. It is also important to note that among the 190 subjects with at least one Tau-PET scan, only 19 of them had one follow-up visit. This means that tau markers dynamics cannot be reliably estimated. Including tau data will require studies on larger cohorts with complete sets of PET imaging acquisitions. This could be part of future extensions of this work, where the inclusion of tau markers will allow to simulate scenarios of production blockage of both amyloid and tau at different rates or intervention time.

Lately, disappointing results of clinical studies led to hypothesize specific treatments targeting AD sub-populations based on their genotype (Safieh et al., 2019). While in our work we describe a global progression of Alzheimer’s disease, in the future we will account for sub-trajectories due to genetic factors, such as the presence of 4 allele of apolipoprotein (APOE4), which is a major risk for developing Alzheimer’s disease influencing both disease onset and progression (Kim et al., 2009). This could be done by estimating dynamical systems specific to the genetic condition of each patient. This was not possible in this study due to a strong imbalance between the number of carriers and non-carriers across the different clinical groups (*cf*. Table 1). Indeed, we observe that the number of ADNI non-carriers is much lower than the number of carriers, especially in the latest stages of the disease (MCI converters and AD). On the contrary, the majority of NL stable subjects are non-carriers. Therefore, applying the model in such conditions would lead to a bias towards more represented groups during the different stages of the disease progression (APOE4-at early stages and APOE4+ at late ones), thus preventing us from differentiating the biomarkers dynamics based on the genetic status. Yet, simulating dynamical relationships specific to genetic factors is a crucial avenue of improvement of our approach, as it would allow to evaluate the effect of APOE4 on intervention time or drug dosage. In addition to this example, there exist numerous non-genetic aggravating factors that may also affect disease evolution, such as diabetes, obesity or smoking. Extending our model to account for panels of risk factors would ultimately allow to test in silico personalized intervention strategies. Moreover, a key aspect of clinical trials is their economic cost. Our model could be extended to help designing clinical trials by optimizing intervention with respect to the available funding. Given a budget, we could simulate scenarios based on different sample size, and trials duration, while estimating the expected cognitive outcome.

Results presented in this work are based on a model estimated by relying solely on a subset of subjects and measures from the ADNI cohort, and therefore they may not be fully representative of the general Alzheimer’s disease progression. Indeed, subjects included in this cohort were either amyloid-positive at baseline, or became amyloid-positive during their follow-up visits. This was motivated by the consideration that evidence of pathological amyloid levels is a necessary condition for diagnosing AD as it puts subjects within the “Alzheimer’s disease continuum” (Jack et al., 2018). By narrowing the list of subjects to a subgroup of amyloid positive we increase the chances of selecting a set of patients likely to develop the disease. Moreover, the inclusion of subjects at various clinical stages allows to span the entire spectrum of morphological and physiological changes affecting the brain. Through the joint analysis of markers of amyloid, neurodegeneration and cognition, our model estimates the average trajectory that best describes the progression of the observed measures when going from NL individuals towards AD patients. The selection of amyloid positive patients aims at increasing the signal of Alzheimer’s pathological changes within this cohort, in order to estimate long-term dynamics for the biomarkers that can be associated to the disease. We believe that this modeling choice is based on a clinically plausible rationale, and allows us to perform our study on a sufficiently large cohort enabling the estimation of our model. Bearing this in mind, we acknowledge the potential presence of bias towards the specific inclusion criterion adopted in this work. Indeed, the present results may provide a limited representation of the pathological temporal window captured by the model. For example, applying the model on a cohort containing amyloid-negative subjects may provide additional insights on the overall disease history. However, this is a challenging task as it would require to identify sub-trajectories dissociated from normal ageing (Lorenzi et al., 2015; Sivera et al., 2020). Another potential bias affecting the results may come from the choice of the clinical scores used to estimate our model. In this study, we relied on a panel of 7 neuro-psychological assessments providing a comprehensive representation of cognitive, memory and functional abilities: ADAS11, MMSE, RAVLT immediate, RAVLT learning, RAVLT forgetting, FAQ, and CDRSB. The choice of these particular scores is consistent with previous literature on DPM (Donohue et al., 2014; Lorenzi et al., 2017). However, it is important to note that our model can handle any type of clinical assessment. Therefore, investigating the effect of adding supplementary clinical scores on the model’s findings would be an interesting future application of our approach, and could be done without any modification of its current formulation. Finally, in addition to these specific characteristics of the cohort, there exists additional biases impacting the model estimation. For instance, the fact that gray matter atrophy and glucose metabolism become abnormal approximately at the same time in Figure 3 can be explained by the high atrophy rate of change in some key regions in normal elders, such as in the hippocampus, compared to the rate of change of FDG (see Table 1). We note that this stronger change of atrophy with respect to glucose metabolism can already be appreciated in the clinically healthy group.

## Conclusion

In this study we investigated a novel quantitative instrument for the development of intervention strategies for disease modifying drugs in AD. Our framework enables the simulation of the effect of intervention time and drug dosage on the evolution of imaging and clinical biomarkers in clinical trials. The proposed data-driven approach is based on the modeling of the spatio-temporal dynamics governing the joint evolution of imaging and clinical measurements throughout the disease. The model is formulated within a Bayesian framework, where the latent representation and dynamics are efficiently estimated through stochastic variational inference. To generate hypothetical scenarios of amyloid lowering interventions, we applied our approach to multi-modal imaging and clinical data from ADNI. The results quantify the crucial role of intervention time, and provide a theoretical justification for testing amyloid modifying drugs in the pre-clinical stage. Our experimental simulations are compatible with the outcomes observed in past clinical trials and suggest that anti-amyloid treatments should be administered at least 7 years earlier than what is currently being done in order to obtain statistically powered improvement of clinical endpoints.

## Abbreviations

DPM: Disease Progression Model
ODE: Ordinary Differential Equations
ADNI: Alzheimer’s Disease Neuroimaging Initiative
NL: Healthy
MCI: Mild Cognitive Impairment
AD: Alzheimer’s dementia
AV45: (18)F-florbetapir Amyloid
FDG: (18)F-fluorodeoxyglucose
ADAS11: Alzheimer’s Disease Assessment Scale
MMSE: Mini-Mental State Examination
FAQ: Functional Assessment Questionnaire
RAVLT: Rey Auditory Verbal Learning Test
CDRSB: Clinical Dementia Rating Scale Sum of Boxes

## Acknowledgements

Data collection and sharing for this project was funded by the Alzheimer’s Disease Neuroimaging Initiative (ADNI) and DOD ADNI. ADNI is funded by the National Institute on Aging, the National Institute of Biomedical Imaging and Bioengineering, and through generous contributions from the following: AbbVie, Alzheimer’s Association; Alzheimer’s Drug Discovery Foundation; Araclon Biotech; BioClinica, Inc.; Biogen; Bristol-Myers Squibb Company;CereSpir, Inc.;Cogstate;Eisai Inc.; Elan Pharmaceuticals, Inc.; Eli Lilly and Company; EuroImmun; F. Hoffmann-La Roche Ltd and its affiliated company Genentech, Inc.; Fujirebio; GE Healthcare; IXICO Ltd.; Janssen Alzheimer Immunotherapy Research & Development, LLC.; Johnson\& Johnson Pharmaceutical Research & Development LLC.;Lumosity;Lundbeck;Merck & Co., Inc.; Meso Scale Diagnostics, LLC.;NeuroRx Research; Neurotrack Technologies;Novartis Pharmaceuticals Corporation; Pfizer Inc.; Piramal Imaging;Servier; Takeda Pharmaceutical Company; and Transition Therapeutics.The Canadian Institutes of Health Research is providing funds to support ADNI clinical sites in Canada. Private sector contributions are facilitated by the Foundation for the National Institutes of Health (www.fnih.org). The grantee organization is the Northern California Institute for Research and Education, and the study is coordinated by the Alzheimer’s Therapeutic Research Institute at the University of Southern California. ADNI data are disseminated by the Laboratory for Neuro Imaging at the University of Southern California.

