## Supplementary for "Simulating the outcome of amyloid treatments in Alzheimer’s disease from imaging and clinical data"

**Supplementary Material**

*Variational inference*

We rewrite $p(\mathbf{X}_i(t)|\mathbf{z}_i(t), \boldsymbol{\sigma}^2, \boldsymbol{\psi})$ as $p(\mathbf{X}_i(t)|\mathbf{z}_i(t_0), \theta_{ODE}, \boldsymbol{\sigma}^2, \boldsymbol{\psi})$. Assuming independence between subjects, the marginal log-likelihood writes as:

$$\begin{aligned}\mathcal{L} &= \sum_i^N \log\left[p(\mathbf{X}_i(t)|\theta_{ODE}, \boldsymbol{\sigma}^2, \boldsymbol{\psi})\right] \\ &= \sum_i^N \log\Big[\int p(\mathbf{X}_i(t)|\mathbf{z}_i(t_0), \theta_{ODE}, \boldsymbol{\sigma}^2, \boldsymbol{\psi})p(\mathbf{z}_i(t_0))d\mathbf{z}_i(t_0)\Big].\end{aligned} \tag{1}$$

For ease of notation, we drop the $i$ index, and dependence on $t$ and $t_0$ is made implicit. Within a Bayesian framework, we wish to maximize $\mathcal{L}$ in order to obtain a posterior distribution for the latent variable $\mathbf{z}$. Since derivation of this quantity is generally not tractable, we resort to stochastic variational inference to tackle the optimization problem. We assume a $\mathcal{N}(\mathbf{0}, \mathbf{I})$ prior for $p(\mathbf{z})$, and introduce an approximate posterior distribution $q(\mathbf{z}|\mathbf{X})$ [2], in order to derive a lower-bound (ELBO) $\mathcal{E}$ for the marginal log-likelihood:

$$\begin{aligned}\log p(\mathbf{X}|\theta_{ODE}, \boldsymbol{\sigma}^2, \boldsymbol{\psi}) &\geq \mathbb{E}_{q(\mathbf{z}|\mathbf{X})}\Big[\log p(\mathbf{X}|\mathbf{z}, \theta_{ODE}, \boldsymbol{\sigma}^2, \boldsymbol{\psi})\Big] \\ &\quad - \mathcal{D}\Big[q(\mathbf{z}|\mathbf{X})|p(\mathbf{z})\Big] \\ &= \mathcal{E},\end{aligned} \tag{2}$$

where $\mathcal{D}$ refers to the Kullback-Leibler (KL) divergence. We propose to factorize the distribution $q(\mathbf{z}|\mathbf{X})$ across modalities such that, $q(\mathbf{z}|\mathbf{X}) = \prod_m q(z^m|\mathbf{x}^m)$, where $q(z^m|\mathbf{x}^m) = \mathcal{N}(f(\mathbf{x}^m, \phi_m^1), h(\mathbf{x}^m, \phi_m^2))$, is a variational Gaussian approximation with moments parameterized by the functions $f$ and $h$. This modality-wise encoding of the data enables to interpret each coordinate of $\mathbf{z}$ as a compressed representation of the corresponding modality. Moreover, the lower-bound simplifies as:

$$\mathcal{E} = \sum_m \mathbb{E}_{q(\mathbf{z}|\mathbf{X})}\Big[\log p(\mathbf{x}^m|\mathbf{z}, \theta_{ODE}, \sigma_m^2, \psi_m)\Big] - \mathcal{D}\Big[q(z^m|\mathbf{x}^m)|p(z^m)\Big]. \tag{3}$$

Details about the ELBO derivation and the computation of the KL divergence are given in sections *Lower bound* and *KL divergence* of this document. A graphical model of the method is also provided in Supplementary figure 1, while Supplementary algorithm 1 details the steps to compute the ELBO.

*Model optimization*

Using the reparameterization trick [3], we can efficiently sample from the posterior distribution $q(\mathbf{z}(t_0)|\mathbf{X}(t_0))$ to approximate the expectation terms. Moreover, thanks to our choices of priors and approximations the KL terms can be computed in closed-form. In practice, we sample from $q(\mathbf{z}(t_0)|\mathbf{X}(t_0))$ to obtain a latent representation $\mathbf{z}(t_0)$ at baseline, while the follow-up points are estimated by decoding the latent time-series obtained through the integration of the ODEs of Eq 3 (in the main manuscript). The model is trained by computing the total ELBO for all the subjects at all the available time points. The parameters $\boldsymbol{\psi}, \boldsymbol{\phi}^1, \boldsymbol{\phi}^2, \theta_{ODE}, \boldsymbol{\sigma}$ are optimized using gradient descent, which requires to backpropagate through the integration operation.

In order to enable backpropagation through the ODEs integration we need to numerically solve the differential equation using only operations that can be differentiated. In this work, we used the Midpoint method which follows a second order Runge-Kutta scheme. The method consists in evaluating the derivative of the solution at $(t_{i+1} + t_i)/2$, which is the midpoint between $t_i$ at which the correct $\mathbf{z}(t)$ is evaluated, and the following $t_{i+1}$:

$$\begin{aligned}\int_{t_i}^{t_{i+1}} g(\mathbf{z}(x))dx &\approx h \cdot g\Big(\mathbf{z}(\frac{t_i + t_{i+1}}{2})\Big) \\ &\approx h \cdot g\Big(\mathbf{z}(t_i) + \frac{h}{2} g(\mathbf{z}(t_i))\Big), \quad h = t_{i+1} - t_i.\end{aligned} \tag{4}$$

Therefore, solving the system of Equation 3 (in the main manuscript) on the interval $[t_0, ..., t]$ only requires operations that can be differentiated, allowing to compute the derivatives of the ELBO with respect to all the parameters, and to optimize them by gradient descent. Moreover, in order to control the variability of the estimated latent trajectory $\mathbf{z}(t)$ due to the error propagation during integration, we initialized the weights of $\boldsymbol{\phi}^1$ and $\boldsymbol{\phi}^2$ such that the approximate posterior of the latent representation for each modality $m$ at baseline was following a $\mathcal{N}(0, 0.01)$ distribution. Finally, we also tested other ODE solvers such as Runge-Kutta 4, which gave similar results than the Midpoint method with a slower execution time due its more expensive approximation scheme.

Concerning the implementation, we trained the model using the ADAM optimizer [4] with a learning rate of 0.01. The functions $f, h$ and $\mu_m$ were parameterized as linear transformations. The model was implemented in Pytorch [5], and we used the *torchdiffeq* package developed in [6] to backpropagate through the ODE solver.

*Lower bound*

We provide here the detailed derivation to obtain the ELBO of Equation 3.

$$\begin{aligned}\log p(\mathbf{X}|\sigma^2, \boldsymbol{\psi}) &= \log\Big[\int p(\mathbf{X}|\mathbf{z}, \theta_{ODE}, \boldsymbol{\sigma^2}, \boldsymbol{\psi})p(\mathbf{z})d\mathbf{z}\Big] \\ &= \log\Big[\int p(\mathbf{X}|\mathbf{z}, \theta_{ODE}, \boldsymbol{\sigma^2}, \boldsymbol{\psi})p(\mathbf{z})\frac{q(\mathbf{z}|\mathbf{X})}{q(\mathbf{z}|\mathbf{X})}d\mathbf{z}\Big] \\ &= \log\Big[\mathbb{E}_{q(\mathbf{z}|\mathbf{X})}\frac{p(\mathbf{X}|\mathbf{z}, \theta_{ODE}, \boldsymbol{\sigma^2}, \boldsymbol{\psi})p(\mathbf{z})}{q(\mathbf{z}|\mathbf{X})}\Big] \\ &\overset{\text{Jensen}}{\geq} \mathbb{E}_{q(\mathbf{z}|\mathbf{X})}\Big[\log\frac{p(\mathbf{X}|\mathbf{z}, \theta_{ODE}, \boldsymbol{\sigma^2}, \boldsymbol{\psi})p(\mathbf{z})}{q(\mathbf{z}|\mathbf{X})}\Big] \\ &= \mathbb{E}_{q(\mathbf{z}|\mathbf{X})}\Big[\log p(\mathbf{X}|\mathbf{z}, \theta_{ODE}, \boldsymbol{\sigma^2}, \boldsymbol{\psi})\Big] - \mathcal{D}\Big[q(\mathbf{z}|\mathbf{X})|p(\mathbf{z})\Big] \\ &= \mathcal{E}.\end{aligned} \tag{5}$$

Given that:

$$p(\mathbf{X}|\mathbf{z}, \theta_{ODE}, \boldsymbol{\sigma^2}, \boldsymbol{\psi}) = \prod_m p(\mathbf{x}^m|\mathbf{z}, \theta_{ODE}, \sigma_m^2, \psi_m), \; q(\mathbf{z}|\mathbf{X}) = \prod_m q(z^m|\mathbf{X}), \text{ and, } p(\mathbf{z}) = \mathcal{N}(\mathbf{0}, \mathbf{I}).$$

We obtain:

$$\mathcal{E} = \sum_m E_{q(\mathbf{z}|\mathbf{X})}\Big[\log p(\mathbf{x}^m|\mathbf{z}, \theta_{ODE}, \sigma_m^2, \psi_m)\Big] - \mathcal{D}\Big[q(z^m|\mathbf{x}^m)|p(z^m)\Big]. \tag{6}$$

*KL divergence*

We have that:

$$\begin{aligned}q(z^m|\mathbf{X}) &= \mathcal{N}(f(\mathbf{x}^m, \phi_m^1), h(\mathbf{x}^m, \phi_m^2)), \\ p(z^m) &= \mathcal{N}(0, 1).\end{aligned} \tag{7}$$

We use the closed-form formula to calculate the KL divergence between two normal distributions:

$$\begin{aligned}\mathcal{D}\Big[q(\mathbf{z}|\mathbf{X})|p(\mathbf{z})\Big] &= \sum_m \mathcal{D}\Big[q(z^m|\mathbf{x}^m)|p(z^m)\Big] \\ &= \frac{1}{2}\sum_m \Big[-\log(h(\mathbf{x}^m, \phi_m^2)) - 1 + h(\mathbf{x}^m, \phi_m^2) + f(\mathbf{x}^m, \phi_m^1)^2\Big]\end{aligned} \tag{8}$$

*Graphical model*

Supplementary figure 1 below provides the graphical model illustrating the method presented in Section *Materials and methods*.

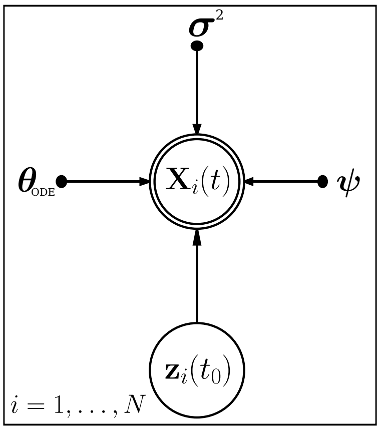


Supplementary figure 1: Graphical model of the proposed method.

*Lower-bound computation*

Supplementary algorithm 1 below details the steps to compute the lower-bound for a given subject $i$ at time $t$.

**Supplementary algorithm 1** Forward pass to compute the lower-bound for a given subject $i$ at time $t$.

**function** COMPUTE_ELBO($\mathbf{X}(t), \mathbf{X}(t_0), \theta_{ODE}, \boldsymbol{\psi}, \boldsymbol{\phi}, \boldsymbol{\sigma}^2$)
For ease of notation we drop the $i$ index in the pseudo-code.
  Sample $\mathbf{z}(t_0) \sim q(\mathbf{z}(t_0)|\mathbf{X}(t_0)) = \prod_m \mathcal{N}(f(\mathbf{x}^m(t_0), \phi_m^1), h(\mathbf{x}^m(t_0), \phi_m^2))$ ▷ Baseline latent representation (reparameterization trick).
  Compute $\mathbf{z}(t) = MIDPOINT(\mathbf{z}(t_0), g, \theta_{ODE}, t)$ ▷ Predict latent representation at time t by numerically solving the ODEs system.
  Compute $\mathbb{E}_{q(\mathbf{z}(t_0)|\mathbf{X}(t_0))}\Big[\log p(\mathbf{x}^m(t)|\mathbf{z}(t), \theta_{ODE}, \sigma_m^2, \psi_m)\Big] \approx -\frac{D_m}{2}\log(2\pi\sigma_m^2) - \frac{1}{2\sigma_m^2}||\mathbf{x}^m(t) - \mu_m(\mathbf{z}(t))||^2$ ▷ Expectation term Equation 3.
  Compute $\mathcal{D}\Big[q(z^m(t_0)|\mathbf{x}^m(t_0))|p(z^m(t_0))\Big] = \frac{1}{2}\Big[-\log(h(\mathbf{x}^m(t_0), \phi_m^2)) - 1 + h(\mathbf{x}^m(t_0), \phi_m^2) + f(\mathbf{x}^m(t_0), \phi_m^1)^2\Big]$ ▷ KL divergence Equation 3.
  Compute $\mathcal{E} = \sum_m \mathbb{E}_{q(\mathbf{z}(t_0)|\mathbf{X}(t_0))}\Big[\log p(\mathbf{x}^m(t)|\mathbf{z}(t), \theta_{ODE}, \sigma_m^2, \psi_m)\Big] - \mathcal{D}\Big[q(z^m(t_0)|\mathbf{x}^m(t_0))|p(z^m(t_0))\Big]$.
  **Return** $\mathcal{E}$
**end function**

*Time-shift comparison and validation*

We compared our estimated disease severity (Figure 2 panel II right in the manuscript) with the one obtained applying the monotonic Gaussian Process (GP) model of (Lorenzi et al., 2017) from the state-of-the-art (Supplementary figure 2A). While both methods estimate significant time differences when going from healthy to pathological stages, our approach captures a larger temporal variability for both earlier and later stages of the disease, as shown in Supplementary figure 2B, highlighting a stronger separability across clinical stages.

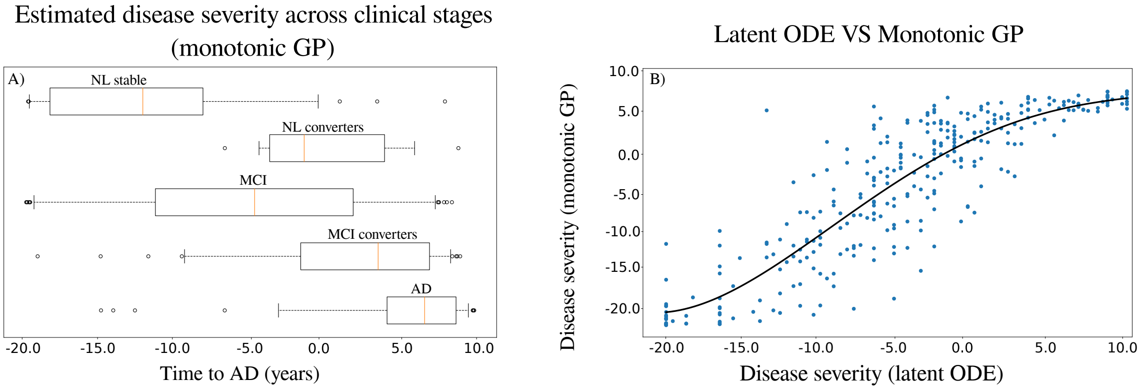


Supplementary figure 2: A: Distribution of the disease severity estimated by the monotonic GP method (Lorenzi et al., 2017) on the training set. NL: normal individuals, MCI: mild cognitive impairment, AD: Alzheimer's dementia. B: Comparison of the disease severity estimated by our method (denoted as latent ODE) with respect to the one estimated by the monotonic GP.

We also assessed the model on an independent testing cohort from the ADNI composed of 130 NL stable, 10 NL converters, 125 MCI stable, 7 MCI converters, and 12 AD subjects which were not necessarily amyloid positive. It is important to note that no PET-FDG data was available for these subjects. We provide in Supplementary table 1 socio-demographic and clinical information for the testing cohort across the different clinical groups. Despite the fact that no FDG data was used to estimate the disease severity, we observe in Supplementary figure 3 that the method still exhibits good separating performances between clinical stages, coherently with the clinical status of the testing individuals.

Supplementary table 1: Baseline socio-demographic and clinical information for testing cohort (284 subjects for 2116 data points). Average values, standard deviation in parenthesis. NL: normal individuals, MCI: mild cognitive impairment, AD: Alzheimer's dementia. ADAS11: Alzheimer's Disease Assessment Scale-cognitive subscale, 11 items. AV45: (18)F-florbetapir Amyloid PET imaging. SUVR: Standardized Uptake Value Ratio.

| | NL stable | NL converters | MCI stable | MCI converters | AD |
|---|---|---|---|---|---|
| N | 130 | 10 | 125 | 7 | 12 |
| Age (yrs) | 72 (6) | 74 (8) | 71 (8) | 73 (9) | 78 (6) |
| Education (yrs) | 17 (2) | 16 (2) | 16 (3) | 14 (3) | 17 (2) |
| ADAS11 | 5.4 (2.8) | 7.7 (4.1) | 7.8 (3.3) | 14.3 (5.2) | 15.0 (6.7) |
| WholeBrain ($cm^3$) | 1063 (103) | 1104 (98) | 1054 (97) | 966 (104) | 1010 (108) |
| AV45 (SUVR) | 0.9 (0.1) | 1.0 (0.1) | 1.0 (0.1) | 1.1 (0.2) | 1.2 (0.3) |

*Simulated clinical endpoints*

We provide in Supplementary table 2 the estimated values for each clinical score at predicted conversion time for the normal progression case when performing the simulations presented in Section *Simulating clinical intervention*.

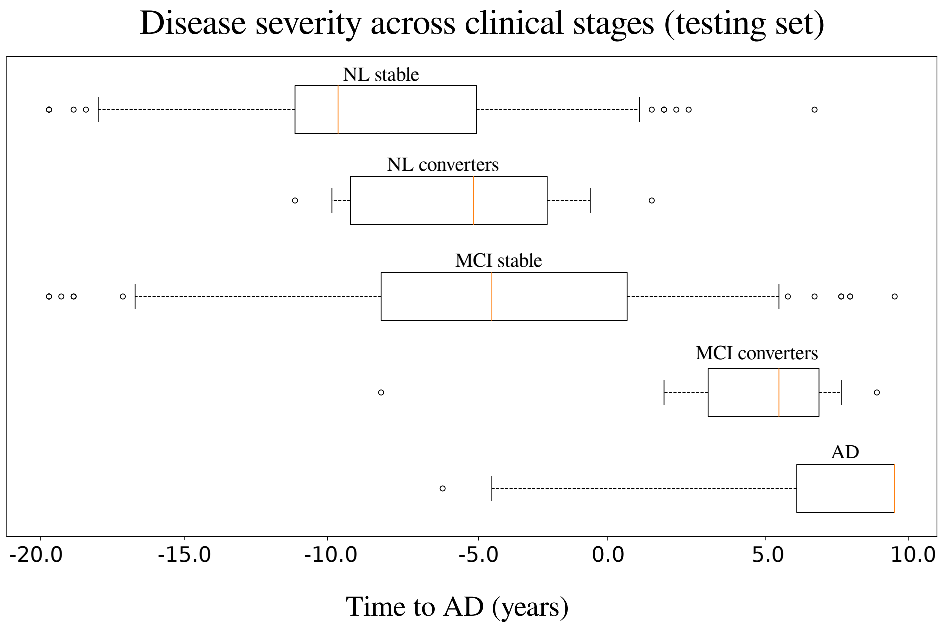


Supplementary figure 3: Distribution of the disease severity estimated for the subjects of the testing set, relatively to the long-term dynamics of Figure 2 panel II) (left) in the manuscript. NL: normal individuals, MCI: mild cognitive impairment, AD: Alzheimer's dementia.

Supplementary table 2: Estimated mean (standard deviation) of the clinical outcomes at predicted conversion time for the normal progression case by year of simulated intervention (100% and 50% amyloid lowering interventions). Results in bold indicate a statistically significant difference between placebo and treated scenarios (p<0.01, two-sided t-test, 100 cases per arm). AD: Alzheimer's dementia, ADAS11: Alzheimer's Disease Assessment Scale, MMSE: Mini-Mental State Examination, FAQ: Functional Assessment Questionnaire, RAVLT: Rey Auditory Verbal Learning Test, CDRSB: Clinical Dementia Rating Scale Sum of Boxes.

**Amyloid lowering intervention 100%**

Score per intervention time

| Score \ Years to AD | −20 | −15 | −12.5 | −10 | −5 | −3 | −2 | −1 |
|---|---|---|---|---|---|---|---|---|
| ADAS11 | **7.8 (8.9)** | **13.7 (5.9)** | **15.9 (5.0)** | **17.3 (4.5)** | 18.6 (4.2) | 18.8 (4.1) | 18.8 (4.1) | 18.9 (4.1) |
| MMSE | **28.3 (3.8)** | **25.7 (2.5)** | **24.8 (2.1)** | **24.2 (2.0)** | 23.6 (1.8) | 23.5 (1.8) | 23.5 (1.8) | 23.5 (1.8) |
| FAQ | **2.3 (7.9)** | **7.5 (5.2)** | **9.3 (4.5)** | **10.5 (4.0)** | 11.7 (3.7) | 11.8 (3.7) | 11.9 (3.7) | 11.9 (3.7) |
| RAVLT immediate | **39.2 (12.7)** | **31.0 (8.4)** | **28.1 (7.1)** | **26.1 (6.4)** | 24.3 (5.8) | 24.0 (5.7) | 23.9 (5.7) | 23.9 (5.7) |
| RAVLT learning | **4.9 (2.2)** | **3.4 (1.5)** | **2.9 (1.2)** | **2.6 (1.1)** | 2.2 (1.0) | 2.2 (1.0) | 2.2 (1.0) | 2.2 (1.0) |
| RAVLT forgetting | **49.9 (29.8)** | **69.4 (19.7)** | **76.6 (16.8)** | **81.3 (15.1)** | 85.9 (13.7) | 86.6 (13.5) | 86.9 (13.4) | 87.0 (13.4) |
| CDRSB | **1.0 (2.8)** | **2.9 (1.9)** | **3.6 (1.6)** | **4.0 (1.5)** | 4.4 (1.4) | 4.5 (1.3) | 4.5 (1.3) | 4.5 (1.3) |

**Amyloid lowering intervention 50%**

Score per intervention time

| Score \ Years to AD | −20 | −15 | −12.5 | −10 | −5 | −3 | −2 | −1 |
|---|---|---|---|---|---|---|---|---|
| ADAS11 | **14.0 (5.9)** | **16.6 (4.8)** | **17.6 (4.5)** | 18.3 (4.4) | 18.9 (4.2) | 19.0 (4.2) | 19.0 (4.2) | 19.0 (4.2) |
| MMSE | **25.6 (2.5)** | **24.5 (2.1)** | **24.0 (2.0)** | 23.7 (1.9) | 23.5 (1.8) | 23.4 (1.8) | 23.4 (1.8) | 23.4 (1.8) |
| FAQ | **7.7 (5.2)** | **10.0 (4.3)** | **10.8 (4.1)** | 11.4 (3.9) | 11.9 (3.7) | 12.0 (3.7) | 12.0 (3.7) | 12.0 (3.7) |
| RAVLT immediate | **30.5 (8.3)** | **27.0 (6.8)** | **25.6 (6.4)** | 24.7 (6.1) | 23.8 (5.8) | 23.7 (5.8) | 23.6 (5.8) | 23.6 (5.8) |
| RAVLT learning | **3.3 (1.5)** | **2.7 (1.2)** | **2.5 (1.1)** | 2.3 (1.1) | 2.2 (1.0) | 2.1 (1.0) | 2.1 (1.0) | 2.1 (1.0) |
| RAVLT forgetting | **70.9 (19.5)** | **79.4 (16.1)** | **82.7 (15.0)** | 84.9 (14.3) | 87.1 (13.7) | 87.4 (13.6) | 87.6 (13.6) | 87.6 (13.6) |
| CDRSB | **3.0 (1.9)** | **3.8 (1.6)** | **4.1 (1.5)** | 4.3 (1.4) | 4.5 (1.4) | 4.5 (1.4) | 4.6 (1.4) | 4.6 (1.4) |
